# Peculiar Pigment Pattern and Population Profile of a Poisonous Pufferfish

**DOI:** 10.1101/2025.11.20.687103

**Authors:** Seita Miyazawa, Hiroyuki Doi, Hiroshi Takahashi, Tomoaki Nishiyama, Shuji Shigenobu, Kazuharu Misawa, Kiyoshi Kikuchi, Harumi Sakai

**Affiliations:** D3 Center, The University of Osaka, Osaka, Japan; Osaka Aquarium NIFREL, Osaka, Japan; Department of Applied Aquabiology, National Fisheries University, Shimonoseki, Yamaguchi, Japan; School of Science, Academic Assembly, University of Toyama, Toyama, Japan; National Institute for Basic Biology, Okazaki, Aichi, Japan; Graduate School of Data Science, Yokohama City University, Yokohama, Kanagawa, Japan; RIKEN Center for Advanced Intelligence Project, Chuo-ku, Tokyo, Japan; Fisheries Laboratory, The University of Tokyo, Hamamatsu, Shizuoka, Japan

## Abstract

Pufferfish are well-known for their toxicity, yet they also exhibit a remarkable diversity of pigment patterns. Mushifugu (*Takifugu exascurus*) is a pufferfish endemic to Japan’s coastal waters and is characterized by conspicuous labyrinthine patterns. Despite being recorded along both the Sea of Japan and the Pacific coasts, it has a limited distribution and is infrequently observed. Aside from its unique body pattern, mushifugu shows little to no morphological differences from other *Takifugu* species, often leading to speculation that it may be an interspecific hybrid. In addition, previous theoretical and empirical studies have shown that complex camouflage-like labyrinthine patterns can emerge through the ‘pattern blending’ caused by hybridization between spotted species, providing support for this possibility. Here, we investigate the phylogenetic origin of mushifugu and its distinctive pattern through population structure analysis and demographic inference in comparison with its closest spotted relative, komonfugu (*T. flavipterus*). Mitochondrial DNA (mtDNA) analysis revealed two regional haplogroups within mushifugu—one in the Sea of Japan (SJ) and the other in the Pacific Ocean (PO). In the haplotype network, the SJ haplogroup formed a distinct cluster, whereas the PO haplogroup appeared as its own cluster connected to the komonfugu haplogroup. By contrast, genome-wide SNP analyses indicated limited structure between the SJ and PO mushifugu populations, while clearly separating mushifugu from komonfugu. Coalescent-based demographic inference suggested that the two species diverged following a bottleneck event in the early Pleistocene. These results confirm that mushifugu is a distinct species rather than a recent interspecific hybrid. Nevertheless, evidence of introgression was detected in both mitochondrial and nuclear genomes, suggesting multiple episodes of past hybridization between mushifugu and komonfugu, highlighting the potentially complex evolutionary processes shaping *Takifugu* species and their pigment patterns.

## Introduction

The genus *Takifugu* consists of pufferfish species primarily distributed in the coastal waters of East Asia, with approximately 25 species currently recognized (Matsuura, 2017). This genus includes torafugu (tiger puffer, *T. rubripes*), which has one of the most compact genomes among vertebrates and was among the first species to have its draft genome sequenced in the early stages of genomic research (Aparicio et al., 2002), as well as other commercially important species such as shōsaifugu (*T. snyderi*) and gomafugu (*T. stictonotus*). Although these pufferfish contain a highly lethal neurotoxin, tetrodotoxin, in their internal organs and/or skin, their exceptional taste has made them highly valued as a food source, particularly in Japan, since ancient times (Sakai, 2021).

Mushifugu (*T. exascurus*) is a small pufferfish species characterized by distinctive labyrinth-like patterns along its flanks (Fig. 1A, B) (Jordan & Snyder, 1901). Despite its conspicuous markings, mushifugu is rarely observed and has a limited distribution. Reports exist from several coastal areas along the Sea of Japan, including Tobishima Island (Yamagata Prefecture), Sado Island (Niigata Prefecture), Kasumi (Hyogo Prefecture), and Tsunoshima (Yamaguchi Prefecture), as well as from the Pacific coast of Japan, specifically Misaki (Kanagawa Prefecture) and Minamiise (Mie Prefecture) (Fujita, 1962; Fujita & Honma, 1991). Owing to limited toxicity data, this species is not approved for human consumption in Japan (Pharmaceutical and Food Safety Bureau, Ministry of Health, Labour and Welfare, 2013; Tatsuno et al., 2021). Furthermore, its ecological traits, particularly reproductive habitats and behavior, remain largely unknown (Matsuura, 2017).

**Figure 1:**
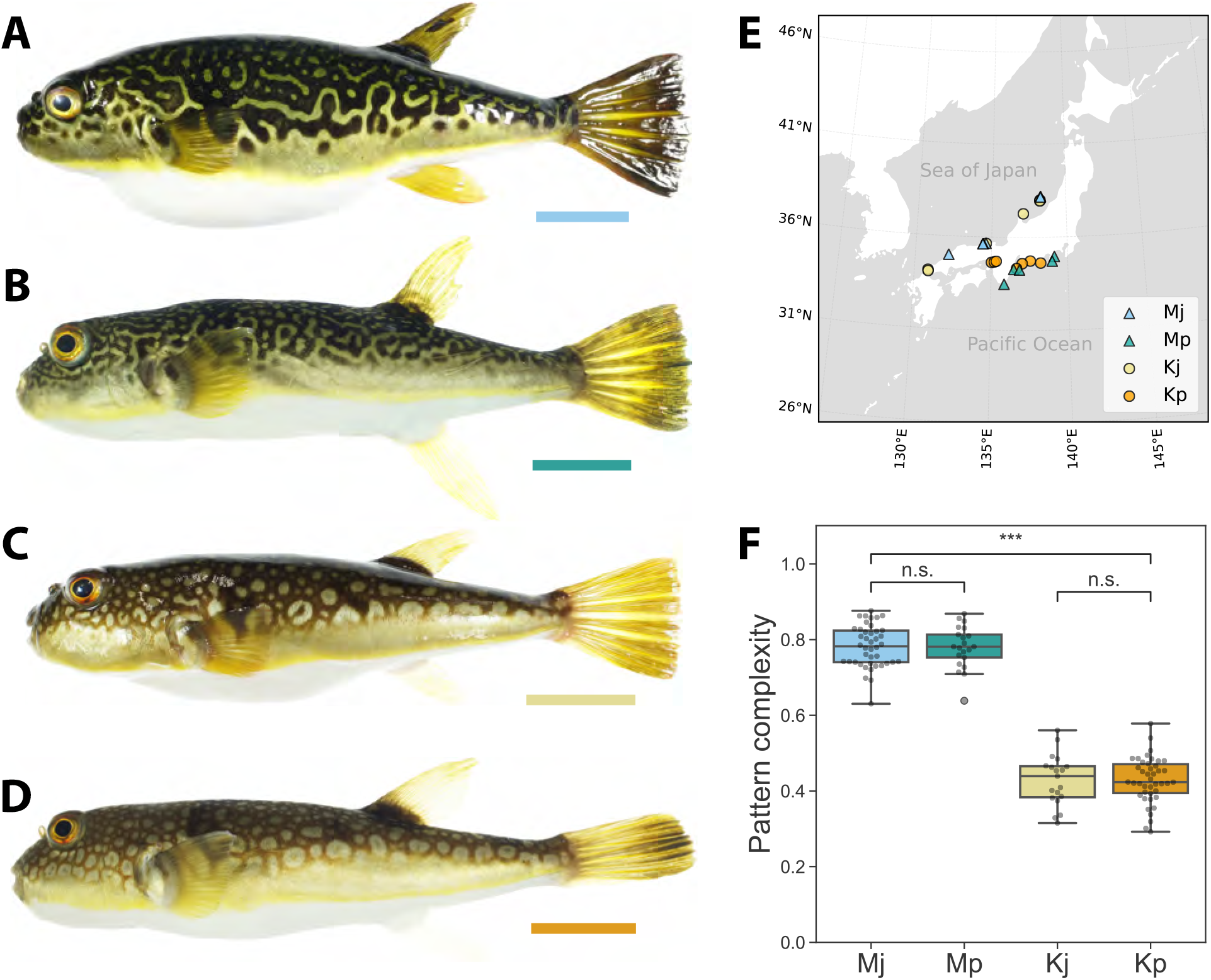
Pigment patterns of pufferfish. Skin pigmentation patterns of mushifugu (*Takifugu exascurus*) from the Sea of Japan (A) and the Pacific coast (B), and komonfugu (*T. flavipterus*, formerly *T. poecilonotus*) from the Sea of Japan (C) and the Pacific coast (D). Sampling locations are shown in (E). (F) Pigment patterns were quantified using the Pattern Complexity Score based on element-wise circularity (Miyazawa, 2020). *** indicates p < 0.001; n.s., not significant. Abbreviations: Mj, mushifugu (Sea of Japan); Mp, mushifugu (Pacific coast); Kj, komonfugu (Sea of Japan); Kp, komonfugu (Pacific coast). Scale bars: 3 cm.

Apart from its distinctive body and caudal-fin pigment pattern, mushifugu exhibits few morphological differences from other species of the genus *Takifugu* (Nakabo, 2013). For example, the fin ray counts of the dorsal, anal, and pectoral fins, as well as the distribution patterns of small spines on the dorsal and ventral surfaces, are identical to those of the white-spotted species, komonfugu (*T. flavipterus*; previously referred to as *T. poecilonotus*) (Fig. 1C, D) (Matsuura, 2017). Given these morphological similarities, its limited collection records, and the known prevalence of interspecific hybridization among *Takifugu* species (Masuda et al., 1991; Miyaki, 1998; Yokogawa & Urayama, 2000; Takahashi et al., 2017), mushifugu has occasionally been suggested to be a hybrid between certain *Takifugu* species rather than a valid species (Nakabo et al., 2012).

Recent theoretical studies on pattern formation suggest that complex, camouflage-like labyrinthine patterns can arise through ‘pattern blending’ between a white-spotted and a black-spotted species as a result of interspecific hybridization (Miyazawa, Watanabe & Kondo, 2021). This prediction has been empirically supported by data from artificial hybridization experiments among salmonid fishes (Miyazawa, Okamoto & Kondo, 2010). Moreover, comparative genomic analyses of the genus *Arothron*, another pufferfish group within the family Tetraodontidae, have provided additional evidence for pattern blending: these studies identified two labyrinthine-patterned species, *Arothron carduus* and *A. multilineatus*, as interspecific hybrids resulting from crosses between the white-spotted species (*A. reticularis*) and the black-spotted species (*A. stellatus*) (Miyazawa, 2020).

*Takifugu* species exhibit a diverse array of patterns, including white spots (e.g., komonfugu, kusafugu (*T. alboplumbeus*; previously referred to as *T. niphobles*), nashifugu (*T. vermicularis*)) and black spots [e.g., higanfugu (*T. pardaris*), gomafugu, shōsaifugu] (Matsuura, 2017), which could serve as potential sources for pattern blending. A previous phylogenetic study using mitochondrial DNA (mtDNA) suggests that *Takifugu* species experienced rapid radiation between 2.4 and 4.7 million years ago (MYA), a timescale comparable to the diversification of cichlids in Lake Malawi (2.4–4.6 MYA) (Yamanoue et al., 2009; Santini et al., 2013). Notably, the white-spotted komonfugu has been identified as the closest known relative of the labyrinthine-patterned mushifugu, with an extremely small—if any—genetic distance observed between them (Yamanoue et al., 2009; Takahashi, Kakioka & Nagano, 2023).

In this study, we aim to elucidate the evolutionary origins of mushifugu and its labyrinthine pattern. To this end, we performed population structure analyses and demographic inference based on mtDNA and whole-genome SNP data.

## Materials & Methods

### Sample collection, body pattern quantification and DNA extraction

A total of 75 individual/tissue specimens of *T. exascurus* and 71 individual/tissue specimens of *T. flavipterus* were collected from various sources (Table S1). The specimens were obtained mainly through fishing along the coasts of the Sea of Japan and the Pacific coast, including the Seto Inland Sea, as well as through purchases from domestic fishery suppliers (Fig. 1E). Additional specimens were donated by fish markets and researchers. Species identification and nomenclature followed Jordan & Snyder (1901) and Matsuura (2017). Body color patterns were quantitatively analyzed in individuals with full-body photographs (Figs. S1 and S2). We focused on the Pattern Complexity Score (PCS), which is defined based on the area-weighted mean isoperimetric quotient of the contours extracted from each image

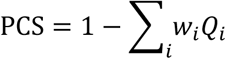

where 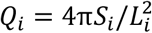 is the isoperimetric quotient (or circularity) of each contour, *w*_*i*_ = *S*_*i*_/ Σ_*i*_ *S*_*i*_ is the area weight, and *S*_*i*_ and *L*_*i*_ are the area and perimeter of each contour, respectively (Miyazawa, 2020). Total DNA was extracted from tissue samples of the pectoral fin and/or skeletal muscle using the DNeasy Blood & Tissue Kit (Qiagen). All animal experiments were conducted in accordance with the institutional guidelines of the University of Osaka.

### Analysis of the mtDNA D-loop Region

The D-loop region of mtDNA (approximately 820–830 bp) was PCR-amplified using KOD-Plus-Neo (TOYOBO) and specific primer sets: forward primer (5’-CTT CCT GAT CCT GAT GCC AAT AG-3’) and reverse primer (5’-TGC GGA TAC TTG CAT GTG TAA GT-3’). The PCR reaction was conducted in a total volume of 20 μL under the following thermal cycling conditions: an initial denaturation at 94°C for 2 min, followed by 35 cycles of denaturation at 98°C for 10 s, annealing at 63°C for 10 s, and extension at 68°C for 1 min, with a final extension at 68°C for 2 min. The PCR products were verified by electrophoresis on a 1% agarose gel and directly sequenced following enzymatic cleanup with ExoSAP-IT (Thermo Fisher Scientific). Sequencing was performed bidirectionally using an Applied Biosystems 3130xl Genetic Analyzer. Multiple alignment of the mtDNA D-loop sequences was performed using MAFFT v7.511 (L-INS-i, default parameters) (Katoh, Rozewicki & Yamada, 2019). The aligned sequences were then used to construct a haplotype network using the median-joining algorithm implemented in the PopART software v1.7 (Leigh & Bryant, 2015) with default parameters.

### Whole genome sequencing and variant calling

Whole genome sequencing was performed for 28 individuals using the Illumina platform. DNA libraries were prepared using the TruSeq DNA PCR-Free Library Prep Kit (Illumina) or the NEBNext Ultra II DNA Library Prep Kit (New England Biolabs) following the manufacturer’s instructions. For one individual each of mushifugu and komonfugu, a series of mate-pair libraries was prepared using the Nextera Mate Pair Library Preparation Kit (Illumina). Sequencing was conducted on the Illumina HiSeq 2000 or HiSeq X platforms. Reads were trimmed with Trimmomatic v0.36 (Bolger, Lohse & Usadel, 2014) and mapped to the reference genome of tiger puffer (*T. rubripes*, assembly fTakRub1.2) using bwa 0.7.17-r1188 (BWA-MEM) (Li, 2013) with the default options. Aligned BAM files were processed using the Genome Analysis Toolkit (GATK, v4.1.7.0) (Van der Auwera & O’Connor, 2020) to mark duplicates (MarkDuplicates) and call variants (HaplotypeCaller). Joint genotyping was performed across all samples using GenotypeGVCFs. Variants were filtered using VariantFiltration with standard hard filtering parameters (QD > 2.0, SOR < 3.0, FS < 60.0, MQ > 40.0, MQRankSum > -12.5 and ReadPosRankSum > -8.0 for SNPs). The filtered variants were used for downstream population structure and demographic analysis.

### Inferring population structure, admixture proportions, and demography

To estimate genetic diversity and differentiation, we calculated genome-wide nucleotide diversity (*π*), absolute divergence (*d*_*XY*_), and *F*_*ST*_ using pixy v2.0.0 (Korunes & Samuk, 2021) from all-sites Variant Call Format (VCF) files. Hudson’s *F*_*ST*_ (Hudson, Slatkin & Maddison, 1992) was derived from genome-wide *π* and *d*_*XY*_, whereas Weir & Cockerham’s *F*_*ST*_ (Weir & Cockerham, 1984) was calculated as a SNP-number–weighted average of estimates obtained in non-overlapping 10 kb windows.

To infer population structure and admixture proportions, we used ADMIXTURE v1.3.0 (Alexander, Novembre & Lange, 2009) on a set of SNPs pruned for linkage desequilibrium (LD) using PLINK v1.90b6.18 (Chang et al., 2015) with the parameters --indep-pairwise 50 10 0.1. Cross-validation was performed for K values ranging from 1 to 5 to determine the most likely number of ancestral populations. Population structure was further assessed through principal component analysis (PCA) using SmartPCA (Price et al., 2006) from the EIGENSOFT package (v7.2.1) (Patterson, Price & Reich, 2006), where the top six principal components were plotted to visualize genetic clustering among individuals.

To test for interspecific introgression, we performed ABBA-BABA tests (D-statistics) using Dsuite (v0.5 r58) (Malinsky, Matschiner & Svardal, 2021) on genome-wide SNPs. For the outgroup, we employed the *Takifugu rubripes* reference genome, the same assembly used for mapping. Significance of excess allele sharing was assessed with block jackknife resampling, and results were summarized as D values, Z-scores, and p-values.

Demographic inference was performed using MSMC2 (v2.1.2) (Schiffels & Wang, 2020) to estimate effective population sizes and the divergence time between the two species from genome-wide SNP data. For population size estimation, each individual was analyzed independently, with two haplotypes per sample. A mutation rate of 5.97×10^−9^ per generation was applied, based on the fish average reported by (Bergeron et al., 2023). The generation time was assumed to be 2.5 years, aligning with estimated maturity ages of 2 years for males and 3 years for females.

Relative cross-coalescence rates (rCCR) between the two species were computed for all 196 pairwise combinations among 14 individuals from each species. For rCCR calculation, phasing was performed with WhatsHap (v2.3) (Martin et al., 2016) using mate-pair and/or paired-end reads. The divergence time between species was estimated as the point at which rCCR reaches a value of 0.5 (Schiffels & Wang, 2020). All results were visualized using a custom Python script.

## Results

### Observation of pigment pattern variations

The observed mushifugu individuals exhibited characteristic labyrinthine body patterns (Figs. 1A, B and S1), clearly distinct from those of all other species in the genus *Takifugu*, including the closely related komonfugu, which displays white-spotted patterns (Figs. 1C, D and S2). Although the specific details of the pattern varied among mushifugu individuals, with differences in the shapes and positions of wavy stripes and occasional spots, each individual consistently exhibited a recognizable maze-like or vermiculated pattern (Fig. S1). This made identification as mushifugu straightforward, in accordance with the diagnostic criteria proposed by Matsuura (Matsuura, 2017). Quantitative analysis of body pattern complexity using the Pattern Complexity Score (PCS) further supported the distinction between mushifugu and komonfugu, showing that the labyrinthine patterns of mushifugu were more complex than the simple spotted patterns of komonfugu (Welch’s *t*-test, *t* = 33.418, *df* = 119.4, *p* = 1.96×10^−62^; mean difference = 0.353, 95% CI 0.332–0.374; Fig. 1F). Within each species, PCS was similar between the Sea of Japan and Pacific populations (Mj vs Mp: *t* = 0.157, *df* = 40.0, *p* = 0.876; Kj vs Kp: *t* = 0.006, *df* = 31.6, *p* = 0.995).

### Analysis of the mtDNA D-loop region and haplotype network construction

Given this phenotypic contrast between labyrinthine-patterned mushifugu and white-spotted komonfugu, we next assessed genetic differentiation by analyzing variation in the mitochondrial control region (D-loop). We examined 58 mushifugu (8 locations) and 55 komonfugu (10 locations) and summarized haplotype relationships in a median-joining network (Fig. 2).

**Figure 2:**
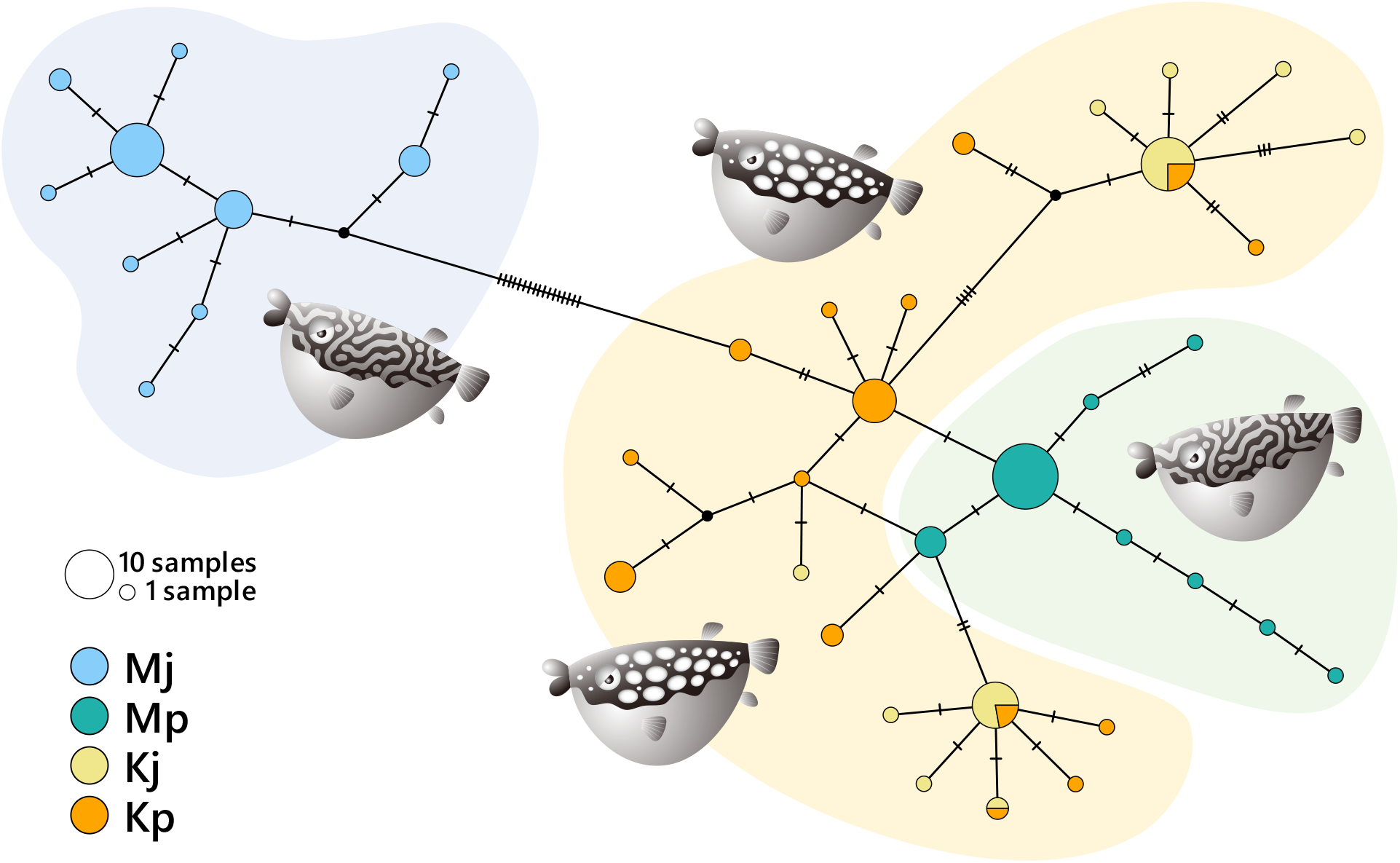
Haplotype network of the mtDNA D-loop region of pufferfish. Haplotype network constructed using the median-joining algorithm based on D-loop sequences from mushifugu and komonfugu. Circle sizes are proportional to haplotype frequencies, and each line denotes a single mutational step. Colors indicate species and sampling locations.

Within mushifugu, we identified two major haplogroups corresponding to populations from the Sea of Japan and the Pacific coast, respectively. Notably, the Pacific mushifugu haplogroup clustered with the komonfugu haplogroup, whereas the Sea of Japan mushifugu haplogroup remained distinct and occurred exclusively within the Sea of Japan population. In contrast, komonfugu exhibited no clear haplogroup differentiation between the Sea of Japan and the Pacific coast populations.

### Population structure analysis based on genome-wide SNPs

To further investigate the genetic relationships between mushifugu and komonfugu, as well as between the Sea of Japan and Pacific populations, we analyzed genome-wide SNP variation from 28 individuals (7 per group: Sea of Japan mushifugu (Mj), Pacific mushifugu (Mp), Sea of Japan komonfugu (Kj), and Pacific komonfugu (Kp)). Whole-genome reads were mapped to the *Takifugu rubripes* reference (fTakRub1.2), with high cross-species mapping rates (97–99%). After quality filtering, a total of 7,396,057 biallelic SNPs were retained. Genome-wide nucleotide diversity (*π*) was lower in mushifugu (≈ 0.0021) than in komonfugu (≈ 0.0032) (Table S2). Genetic differentiation (*F*_*ST*_) was high between mushifugu and komonfugu (≈ 0.41), moderate between the Sea of Japan and Pacific populations of mushifugu (Mj vs. Mp, ≈ 0.20), and very low between those of komonfugu (Kj vs. Kp, ≈ 0.015) (Table S3). After linkage disequilibrium (LD) pruning, 416,407 SNPs were used for subsequent population structure analysis.

Admixture analysis indicated that the optimal number of ancestral populations was K = 2, with K = 3 yielding a comparably low cross-validation error (Fig. 3A, B). At K = 2, the samples were clearly divided into mushifugu and komonfugu, perfectly aligned with the species identification based on pigment patterns. At K = 3, mushifugu was further separated into the Sea of Japan and Pacific populations. At K = 4, an additional weak structure was suggested within komonfugu, roughly aligned with the Sea of Japan and Pacific populations, although the correspondence with geographic locations was not exact.

**Figure 3:**
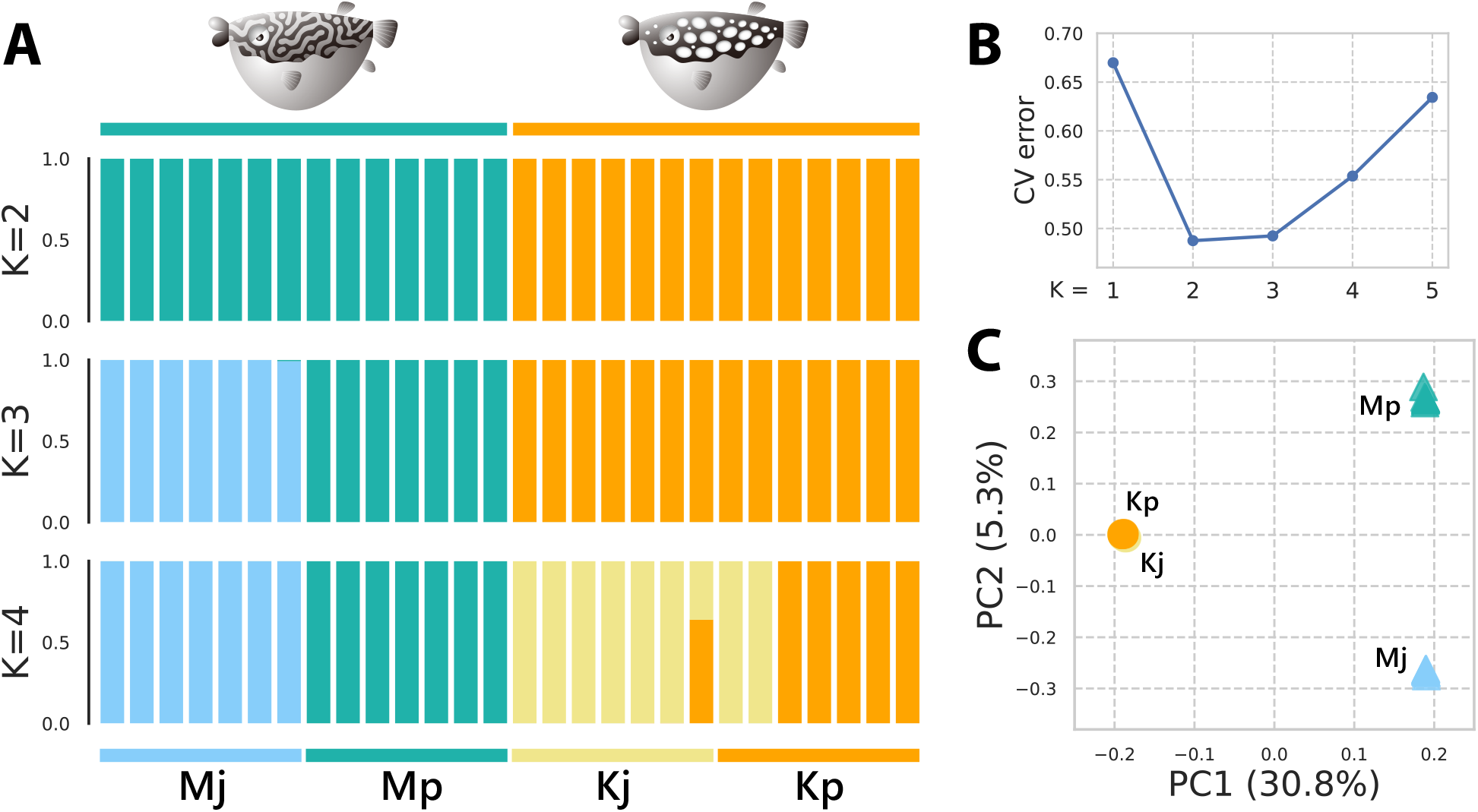
Population structure of mushifugu and komonfugu based on genome-wide SNP data. (A) Admixture analysis at K=2, 3, and 4. (B) Cross-validation error across K values, with the lowest errors at K=2 and 3. (C) Principal component analysis (PCA) of genome-wide SNP variation, where PC1 (30.8%) separates the two species and PC2 (5.3%) reflects geographic variation within mushifugu.

Similar results were obtained from principal component analysis (PCA), where the first principal component (PC1) accounted for 30.8% of the total variance and captured the genetic differentiation between mushifugu and komonfugu (Fig. 3C). The second principal component (PC2) explained 5.3% of the variance and distinguished between the Sea of Japan and Pacific populations within mushifugu. These results indicate a clear genetic differentiation between mushifugu and komonfugu, as well as geographic population divergence within mushifugu. In contrast, within komonfugu, no clear separation between the Sea of Japan and Pacific populations was observed, although the Pacific populations appeared to be more scattered along the third to sixth principal components (Fig. S3).

### Estimation of effective population size and divergence time

We used a coalescent-based approach with genome sequence data to estimate the effective population size and divergence time between mushifugu and komonfugu (Fig. 4). Both species experienced a bottleneck approximately 0.4 to 0.8 MYA in the early Pleistocene, followed by a shared transition pattern of population size increase, and subsequent decline during the Last Glacial Period (Fig. 4A). However, differences were observed in the post-bottleneck trends between the two species: mushifugu exhibited a more moderate expansion compared to komonfugu, and its subsequent population decline was pronounced. These contrasting patterns suggest that the two species gradually diverged following the bottleneck, a conclusion further supported by the analysis of the relative cross-coalescence rate (rCCR; Fig. 4B). The divergence time between the two species was estimated to be approximately 0.26 MYA based on the point at which the rCCR reached 0.5.

**Figure 4:**
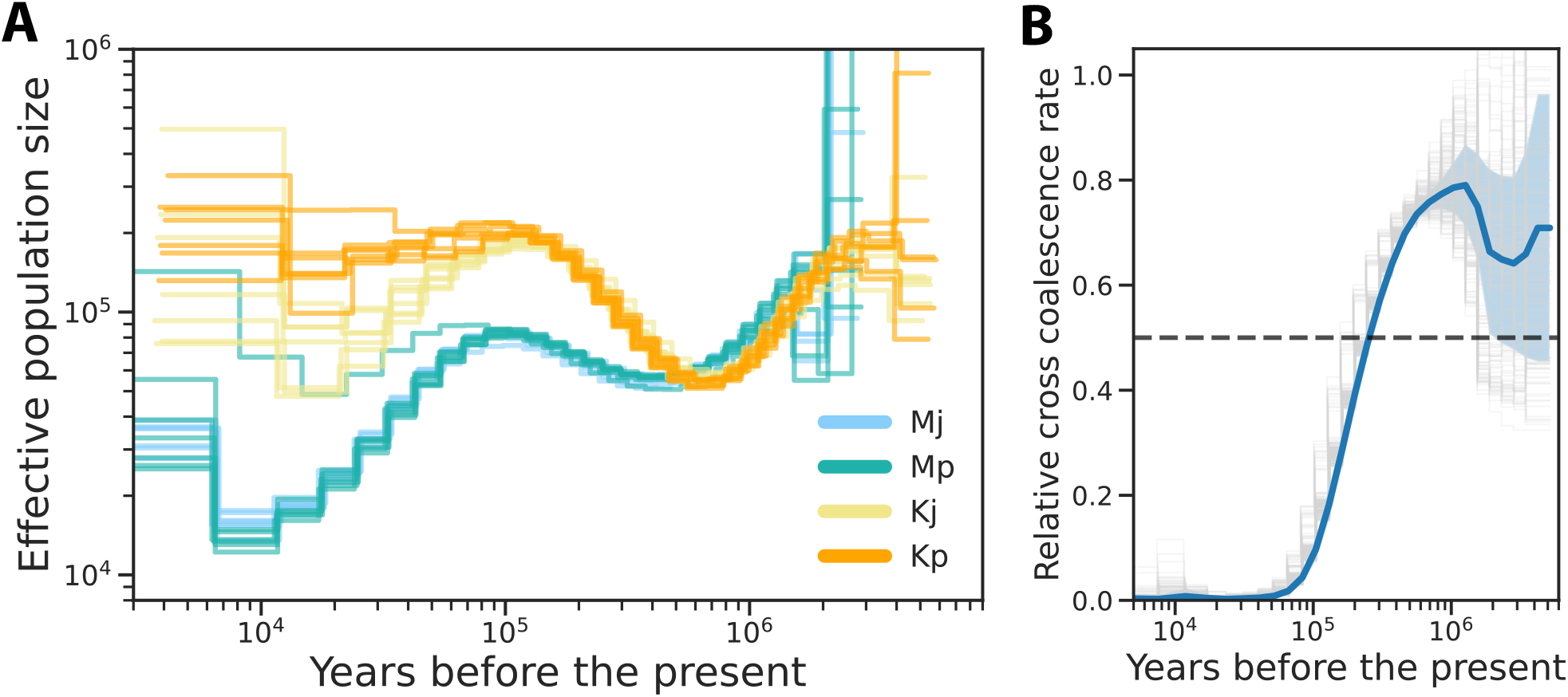
Demographic history of mushifugu and komonfugu inferred from genome-wide SNP data. (A) Effective population sizes inferred using MSMC2, with two haplotypes per individual. A mutation rate of 5.97×10^−9^ per generation (Bergeron et al. 2023) and a generation time of 2.5 years (2 years for males, 3 years for females) were assumed. (B) Relative cross-coalescence rate (rCCR) between the two species, calculated for all 196 pairwise combinations among 14 individuals from each species. The blue solid line shows the mean rCCR, shaded areas indicate ±1 standard deviation, thin gray step lines represent individual pairs, and the dotted line marks rCCR = 0.5.

### Examination of interspecific introgression

Genome-wide data confirmed clear genetic differentiation between mushifugu and komonfugu. In this context, the mtDNA haplotype network, in which Pacific mushifugu clustered more closely with komonfugu than with Sea of Japan mushifugu, suggested interspecific introgression from komonfugu into mushifugu. To further examine this possibility, we conducted ABBA-BABA tests using genome-wide SNP data (Table S4). Significant excess of shared derived alleles was detected between komonfugu and Sea of Japan mushifugu (Mj), with D values ranging from 0.0042 to 0.0075 (Z = 2.0–3.2, p = 0.044–0.0015), indicating introgression from komonfugu into Sea of Japan mushifugu. In contrast, no significant signal was detected for introgression from komonfugu into Pacific mushifugu (Mp), or from mushifugu into komonfugu (Kj or Kp).

## Discussion

In this study, we addressed the evolutionary origin and phylogenetic placement of the pufferfish mushifugu (*T. exascurus*), which exhibits a distinctive labyrinthine pigmentation pattern. Genome-wide evidence indicates strong species-level divergence between mushifugu and its closest spotted relative, komonfugu (*T. flavipterus*).

Previous theoretical and empirical studies have suggested that labyrinthine patterns can emerge through hybridization between spotted species (Miyazawa, Okamoto & Kondo, 2010; Miyazawa, 2020; Miyazawa, Watanabe & Kondo, 2021), and morphological comparisons have raised the possibility that mushifugu may represent interspecific hybrids rather than a valid species (Nakabo et al., 2012). However, our genomic analyses support species status for mushifugu, and demographic inferences from whole-genome data indicate that the two lineages gradually diverged following a bottleneck in the early Pleistocene.

At a finer scale, our population structure analysis revealed two geographically distinct groups within mushifugu: one distributed along the Sea of Japan coast and the other along the Pacific coast. The genetic divergence between these two mushifugu populations was consistently supported by a relatively high *F*_*ST*_ value (≈ 0.2), as well as by ADMIXTURE and PCA analyses (Fig. 3). However, with respect to phenotypic traits such as pattern complexity, no pronounced differences were observed between them (Fig. 1F and S1); thus, their divergence remains cryptic at the morphological level.

Similar cases of genetic differentiation between Sea of Japan and Pacific populations have also been documented in other coastal fishes of the Japanese archipelago, including gobies and sticklebacks (Higuchi & Goto, 1996; Akihito et al., 2008, 2016; Ravinet et al., 2018; Hirase, 2022; Kato et al., 2024). Such differentiation has often been attributed to sea-level fluctuations during the glacial–interglacial cycles of the Pleistocene, particularly the isolation of the Sea of Japan during the Last Glacial Maximum (Hirase, 2022). In mushifugu as well, both the Sea of Japan and Pacific populations experienced reductions in effective population size during the period corresponding to the Last Glacial Maximum (Fig. 4), suggesting that their divergence may likewise be associated with the closure of the Sea of Japan at that time.

Notably, evidence for past hybridization events with komonfugu was detected in both mushifugu populations. Haplotype network analysis of the mitochondrial D-loop region suggested that the Pacific mushifugu population may have experienced interspecific hybridization with komonfugu, resulting in mtDNA introgression. In contrast, genome-wide ABBA-BABA tests indicated significant nuclear introgression from komonfugu into Sea of Japan mushifugu. These seemingly contrasting results are most parsimoniously explained by multiple episodes of interspecific gene flow after the divergence of Pacific and Sea of Japan mushifugu lineages, involving asymmetric introgression of mitochondrial and nuclear genomes.

Interspecific hybridization in the genus *Takifugu* has been documented both under experimental conditions and in the wild (Fujita, 1967; Miyaki, 1992, 1998; Matsuura, 2017). Artificial hybridization experiments have demonstrated that fertile hybrids can be produced even between species with markedly different body sizes and ecological traits, such as torafugu (*T. rubripes*) and kusafugu (*T. alboplumbeus*) (Miyaki, 1998), and that such hybrids can be maintained as interspecific lines for quantitative trait locus (QTL) analysis (Hosoya et al., 2013a,b, 2015; Kim et al., 2019).

In nature, putative hybrid individuals have been observed among various *Takifugu* species, including torafugu, gomafugu (*T. stictinotus*), and shōsaifugu (*T. snyderi*) (Matsuura, 2017). In some cases, the parental species combinations have been clearly identified, such as shimafugu×nashifugu and nashifugu×komonfugu (Masuda et al., 1991; Yokogawa & Urayama, 2000). More recently, a high proportion of wild-caught individuals have been identified as natural hybrids between gomafugu and shōsaifugu (Takahashi et al., 2017).

Our present findings, which suggest post-divergence gene flow between mushifugu and komonfugu, support the notion that interspecific hybridization and genetic introgression are prevalent and ongoing phenomena within the genus *Takifugu* (Takahashi et al., 2017; Liu et al., 2021; Takahashi, 2022).

These findings also raise the possibility that interspecific hybridization and genetic introgression have occurred repeatedly throughout the evolutionary history of *Takifugu* species. Although hybridization has traditionally been viewed as a barrier to species divergence and speciation (Coyne & Orr, 2004), accumulating evidence suggests that it can facilitate speciation and positively contribute to evolutionary processes via genetic introgression (Arnold, 1997, 2016; Mallet, 2007). For example, studies of Darwin’s finches in the Galápagos Islands have demonstrated that interspecific hybridization can promote rapid speciation in favorable ecological contexts (Grant & Grant, 1992; Lamichhaney et al., 2018). Similarly, admixture with archaic hominins such as Neanderthals and Denisovans has been shown to play a significant role in the evolutionary history of modern humans (Green et al., 2010; Huerta-Sánchez et al., 2014; Vernot et al., 2016). It would therefore be intriguing to explore whether genetic introgression has also contributed to the diversification and adaptation of pufferfish lineages.

The genus *Takifugu* comprises species that exhibit a wide range of pigmentation patterns, including spots and stripes. These species are thought to have diversified through explosive speciation events that occurred relatively recently—approximately 2.4–4.7 million years ago—a timescale comparable to the well-known cichlid radiation in Lake Malawi (2.4–4.6 MYA) (Yamanoue et al., 2009; Santini et al., 2013). In African cichlids, such rapid adaptive radiation has been proposed to be facilitated by increased genetic diversity introduced via hybridization (Meier et al., 2017; Malinsky et al., 2018).

Theoretical studies of pattern formation have hypothesized that complex pigmentation patterns can emerge through ‘pattern blending’ resulting from interspecific hybridization (Miyazawa, Okamoto & Kondo, 2010; Miyazawa, 2020). Accordingly, investigating the role of hybridization in the evolution of pigment pattern diversity may yield valuable insights. Future research incorporating other pufferfish species could further elucidate the mystery of how body patterns have evolved and how they may evolve, providing a pivotal piece of the pigmentation puzzle.

## Supplemental Information

Supplementary Table S1–S4 and Supplementary Figures S1–S3.

## Additional Information and Declarations

### Competing Interests

The authors declare there are no competing interests.

### DNA Deposition

The sequencing data have been deposited in DDBJ under BioProject PRJDB35841, BioSample SAMD01606002–SAMD01606029, and Sequence Read Archive (DRA) accession number DRA021836. The mitochondrial D-loop sequences have been deposited in DDBJ under accession numbers LC903457–LC903569.

### Data Availability

The custom code will be available at GitHub (https://github.com/seita42/puffer-popgen) and Zenodo.

### Funding

This work was supported by JST FOREST Program (Grant Number JPMJFR214E) and JSPS KAKENHI (Grant Numbers JP23K27229, JP23H02538, JP15H04415, JP19H03283, and JP25128709).

## Acknowledgements

We thank M. Murata, Y. Sato, S. Awata, H. Ando, T. Shimotani, K. Yamamoto, S. Kuwayama, M. Nagata, K. Kajitani, M. Nishikawa, T. Sonoyama, and Shimonoseki Marine Science Museum Kaikyokan for providing materials; K. Matsuura, Y. Yamanoue, K. Kuriiwa, S. Hosoya, and M. Yamaguchi for valuable discussions and materials; Y. Jo, M. Fujita, K. Yamaguchi, N. Tsukamoto, T. Shibata, and S. Ohi for technical assistance; and M. Hasebe and S. Kondo for their continuous encouragement and support. Computations were partially performed on the NIG supercomputer at ROIS National Institute of Genetics.

## Notes

https://github.com/seita42/puffer-popgen

### Competing Interest Statement

The authors have declared no competing interest.

