## Supplementary Tables S1-S4 and Figures S1-S3 for "Peculiar Pigment Pattern and Population Profile of a Poisonous Pufferfish"

**Table S1: Metadata of *Takifugu exascurus* (mushifugu) and *T. flavipterus* (komonfugu) samples used in this study.**

Details of individual samples analyzed in this study. Columns include sample ID, species, population (SJ: Sea of Japan; PO: Pacific Ocean), collection locality, collection date, availability of body pattern image (1: available, 0: not available), pattern metrics (lightness and complexity), availability of mitochondrial D-loop sequence (1: available, 0: not available), whole-genome resequencing data (NGS; 1: available, 0: not available), and BioSample accession numbers (if assigned).

| ID | Species | Population | Location | Date | Image | Lightness | Complexity | D-loop | NGS | BioSample |
|---|---|---|---|---|---|---|---|---|---|---|
| Mj_002 | Mushifugu | SJ | Sado Island, Niigata Pref. | 2006-06 | 1 | 0.38396 | 0.76110 | 1 | 0 | |
| Mj_003 | Mushifugu | SJ | Sado Island, Niigata Pref. | 2006-06 | 1 | 0.37006 | 0.72597 | 1 | 1 | SAMD01606002 |
| Mj_004 | Mushifugu | SJ | Sado Island, Niigata Pref. | 2006-06 | 1 | 0.37220 | 0.74210 | 1 | 0 | |
| Mj_005 | Mushifugu | SJ | Sado Island, Niigata Pref. | 2006-06 | 1 | 0.32359 | 0.73005 | 1 | 0 | |
| Mj_006 | Mushifugu | SJ | Sado Island, Niigata Pref. | 2006-06 | 1 | 0.36564 | 0.77450 | 1 | 1 | SAMD01606003 |
| Mj_017 | Mushifugu | SJ | Hamasaka, Hyogo Pref. | 2006-04-27 | 1 | 0.39043 | 0.87624 | 1 | 0 | |
| Mj_058 | Mushifugu | SJ | Gotsu, Shimane Pref. | 2007-06-15 | 1 | 0.36045 | 0.73408 | 1 | 0 | |
| Mj_065 | Mushifugu | SJ | Kasumi, Hyogo Pref. | 2012-06-27 | 1 | 0.39179 | 0.74039 | 1 | 1 | SAMD01606004 |
| Mj_074 | Mushifugu | SJ | Hamasaka, Hyogo Pref. | NA | 0 | NA | NA | 1 | 0 | |
| Mj_075 | Mushifugu | SJ | Hamasaka, Hyogo Pref. | NA | 0 | NA | NA | 1 | 0 | |
| Mj_076 | Mushifugu | SJ | Sado Island, Niigata Pref. | NA | 0 | NA | NA | 1 | 0 | |
| Mj_091 | Mushifugu | SJ | Sado Island, Niigata Pref. | 2013-06-05 | 1 | 0.38844 | 0.86359 | 1 | 0 | |
| Mj_092 | Mushifugu | SJ | Sado Island, Niigata Pref. | 2013-06-05 | 1 | 0.38061 | 0.75647 | 1 | 0 | |
| Mj_093 | Mushifugu | SJ | Sado Island, Niigata Pref. | 2013-06-05 | 1 | 0.37332 | 0.82750 | 1 | 0 | |
| Mj_098 | Mushifugu | SJ | Sado Island, Niigata Pref. | 2013-06-05 | 1 | 0.33212 | 0.63052 | 1 | 1 | SAMD01606005 |
| Mj_099 | Mushifugu | SJ | Sado Island, Niigata Pref. | 2013-06-05 | 1 | 0.30407 | 0.69828 | 1 | 0 | |
| Mj_100 | Mushifugu | SJ | Sado Island, Niigata Pref. | 2013-06-05 | 1 | 0.36573 | 0.80173 | 1 | 0 | |
| Mj_101 | Mushifugu | SJ | Sado Island, Niigata Pref. | 2013-06-05 | 1 | 0.36353 | 0.73733 | 1 | 1 | SAMD01606006 |
| Mj_102 | Mushifugu | SJ | Sado Island, Niigata Pref. | 2013-06-05 | 1 | 0.37397 | 0.86274 | 1 | 0 | |
| Mj_103 | Mushifugu | SJ | Sado Island, Niigata Pref. | 2013-06-05 | 1 | 0.33853 | 0.85768 | 1 | 0 | |
| Mj_104 | Mushifugu | SJ | Sado Island, Niigata Pref. | 2013-06-05 | 1 | 0.34003 | 0.78513 | 1 | 0 | |
| Mj_115 | Mushifugu | SJ | Sado Island, Niigata Pref. | 2014-06-11 | 1 | 0.32841 | 0.80248 | 1 | 0 | |
| Mj_116 | Mushifugu | SJ | Sado Island, Niigata Pref. | 2014-06-11 | 1 | 0.38062 | 0.72082 | 1 | 0 | |
| Mj_117 | Mushifugu | SJ | Sado Island, Niigata Pref. | 2014-06-11 | 1 | 0.37864 | 0.74222 | 1 | 1 | SAMD01606007 |
| Mj_118 | Mushifugu | SJ | Sado Island, Niigata Pref. | 2014-06-11 | 1 | 0.33880 | 0.81647 | 1 | 0 | |
| Mj_119 | Mushifugu | SJ | Sado Island, Niigata Pref. | 2014-06-11 | 1 | 0.37093 | 0.82879 | 1 | 0 | |
| Mj_120 | Mushifugu | SJ | Sado Island, Niigata Pref. | 2014-06-11 | 1 | 0.35290 | 0.77955 | 1 | 0 | |
| Mj_121 | Mushifugu | SJ | Sado Island, Niigata Pref. | 2014-06-11 | 1 | 0.36431 | 0.80841 | 0 | 0 | |
| Mj_122 | Mushifugu | SJ | Sado Island, Niigata Pref. | 2014-06-11 | 1 | 0.39210 | 0.81985 | 0 | 0 | |
| Mj_123 | Mushifugu | SJ | Sado Island, Niigata Pref. | 2014-06-11 | 1 | 0.36641 | 0.78789 | 0 | 0 | |
| Mj_124 | Mushifugu | SJ | Sado Island, Niigata Pref. | 2014-06-11 | 1 | 0.36630 | 0.86284 | 0 | 0 | |
| Mj_125 | Mushifugu | SJ | Sado Island, Niigata Pref. | 2014-06-11 | 1 | 0.36592 | 0.79356 | 0 | 0 | |
| Mj_126 | Mushifugu | SJ | Sado Island, Niigata Pref. | 2014-06-11 | 1 | 0.36075 | 0.73700 | 0 | 0 | |
| Mj_127 | Mushifugu | SJ | Sado Island, Niigata Pref. | 2014-06-11 | 1 | 0.36676 | 0.82463 | 0 | 0 | |
| Mj_128 | Mushifugu | SJ | Sado Island, Niigata Pref. | 2014-06-11 | 1 | 0.35058 | 0.77783 | 0 | 0 | |
| Mj_129 | Mushifugu | SJ | Sado Island, Niigata Pref. | 2014-06-11 | 1 | 0.32652 | 0.74030 | 0 | 0 | |
| Mj_130 | Mushifugu | SJ | Sado Island, Niigata Pref. | 2014-06-11 | 1 | 0.34724 | 0.69299 | 0 | 0 | |
| Mj_131 | Mushifugu | SJ | Sado Island, Niigata Pref. | 2014-06-11 | 1 | 0.37541 | 0.83735 | 0 | 0 | |
| Mj_132 | Mushifugu | SJ | Sado Island, Niigata Pref. | 2014-06-11 | 1 | 0.34925 | 0.84632 | 0 | 0 | |
| Mj_133 | Mushifugu | SJ | Sado Island, Niigata Pref. | 2014-06-11 | 1 | 0.34671 | 0.80918 | 0 | 0 | |
| Mj_134 | Mushifugu | SJ | Sado Island, Niigata Pref. | 2014-06-11 | 1 | 0.34762 | 0.75395 | 0 | 0 | |
| Mj_135 | Mushifugu | SJ | Sado Island, Niigata Pref. | 2014-06-11 | 1 | 0.36334 | 0.74283 | 0 | 0 | |
| Mj_163 | Mushifugu | SJ | Sado Island, Niigata Pref. | 2015-06-03 | 1 | 0.34904 | 0.82190 | 1 | 0 | |
| Mj_164 | Mushifugu | SJ | Sado Island, Niigata Pref. | 2015-06-03 | 1 | 0.39551 | 0.85878 | 1 | 1 | SAMD01606008 |
| Mj_165 | Mushifugu | SJ | Sado Island, Niigata Pref. | 2015-06-03 | 1 | 0.33854 | 0.73100 | 1 | 0 | |
| Mp_001 | Mushifugu | PO | Mie Pref. | NA | 1 | 0.34591 | 0.85613 | 1 | 0 | |
| Mp_007 | Mushifugu | PO | Shimoda, Shizuoka Pref. | 2000-06-19 | 0 | NA | NA | 1 | 0 | |
| Mp_008 | Mushifugu | PO | Ito, Shizuoka Pref. | 2000-06-15 | 0 | NA | NA | 1 | 0 | |
| Mp_009 | Mushifugu | PO | Shimoda, Shizuoka Pref. | 2002-09-02 | 0 | NA | NA | 1 | 0 | |
| Mp_010 | Mushifugu | PO | Mie Pref. | 2004-05-01 | 0 | NA | NA | 1 | 0 | |
| Mp_011 | Mushifugu | PO | Minamiise, Mie Pref. | 2004-10-08 | 0 | NA | NA | 1 | 0 | |
| Mp_012 | Mushifugu | PO | Minamiise, Mie Pref. | 2007-02-13 | 1 | 0.38956 | 0.71393 | 1 | 0 | |
| Mp_013 | Mushifugu | PO | Minamiise, Mie Pref. | 2007-02-13 | 1 | 0.40764 | 0.73519 | 1 | 1 | SAMD01606009 |
| Mp_014 | Mushifugu | PO | Minamiise, Mie Pref. | 2007-02-13 | 1 | 0.37515 | 0.77308 | 1 | 1 | SAMD01606010 |
| Mp_015 | Mushifugu | PO | Minamiise, Mie Pref. | 2007-02-13 | 1 | 0.38042 | 0.78133 | 1 | 0 | |
| Mp_059 | Mushifugu | PO | Minamiise, Mie Pref. | 2007-12 | 1 | 0.38173 | 0.81340 | 0 | 0 | |
| Mp_060 | Mushifugu | PO | Minamiise, Mie Pref. | 2007-12 | 1 | 0.33749 | 0.72697 | 0 | 0 | |
| Mp_061 | Mushifugu | PO | Kushimoto, Wakayama Pref. | 2008-05-01 | 0 | NA | NA | 1 | 0 | |
| Mp_062 | Mushifugu | PO | Kushimoto, Wakayama Pref. | 2008-05-01 | 0 | NA | NA | 1 | 1 | SAMD01606011 |

| ID | Species | Population | Location | Date | Image | Lightness | Complexity | D-loop | NGS | BioSample |
|---|---|---|---|---|---|---|---|---|---|---|
| Mp_063 | Mushifugu | PO | Kushimoto, Wakayama Pref. | 2008-06-01 | 0 | NA | NA | 1 | 1 | SAMD01606012 |
| Mp_064 | Mushifugu | PO | Kushimoto, Wakayama Pref. | 2008-06-01 | 0 | NA | NA | 1 | 0 | |
| Mp_114 | Mushifugu | PO | Minamiise, Mie Pref. | 2014-01 | 1 | 0.39784 | 0.80500 | 1 | 1 | SAMD01606013 |
| Mp_178 | Mushifugu | PO | Minamiise, Mie Pref. | 2016-11-07 | 1 | 0.38672 | 0.79041 | 1 | 0 | |
| Mp_204 | Mushifugu | PO | Minamiise, Mie Pref. | 2017-04-13 | 1 | 0.34706 | 0.75273 | 1 | 0 | |
| Mp_205 | Mushifugu | PO | Minamiise, Mie Pref. | 2017-04-13 | 1 | 0.35346 | 0.83292 | 1 | 1 | SAMD01606014 |
| Mp_206 | Mushifugu | PO | Minamiise, Mie Pref. | 2017-04-13 | 1 | 0.30181 | 0.75870 | 1 | 0 | |
| Mp_207 | Mushifugu | PO | Minamiise, Mie Pref. | 2017-04-13 | 1 | 0.35953 | 0.78125 | 1 | 0 | |
| Mp_208 | Mushifugu | PO | Minamiise, Mie Pref. | 2017-04-13 | 1 | 0.33713 | 0.76746 | 1 | 1 | SAMD01606015 |
| Mp_209 | Mushifugu | PO | Minamiise, Mie Pref. | 2017-04-13 | 1 | 0.36613 | 0.77818 | 1 | 0 | |
| Mp_210 | Mushifugu | PO | Minamiise, Mie Pref. | 2017-04-13 | 1 | 0.33626 | 0.63840 | 1 | 0 | |
| Mp_211 | Mushifugu | PO | Minamiise, Mie Pref. | 2017-04-13 | 1 | 0.34798 | 0.84888 | 1 | 0 | |
| Mp_212 | Mushifugu | PO | Minamiise, Mie Pref. | 2017-04-13 | 1 | 0.34801 | 0.70902 | 1 | 0 | |
| Mp_213 | Mushifugu | PO | Minamiise, Mie Pref. | 2017-04-13 | 1 | 0.36483 | 0.80988 | 1 | 0 | |
| Mp_214 | Mushifugu | PO | Minamiise, Mie Pref. | 2017-04-13 | 1 | 0.36761 | 0.83052 | 1 | 0 | |
| Mp_215 | Mushifugu | PO | Shima, Mie Pref. | NA | 1 | 0.45159 | 0.86856 | 1 | 0 | |
| Kj_020 | Komonfugu | SJ | Shimonoseki, Yamaguchi Pref. | 2006-06-18 | 0 | NA | NA | 1 | 1 | SAMD01606016 |
| Kj_021 | Komonfugu | SJ | Shimonoseki, Yamaguchi Pref. | 2006-06-18 | 0 | NA | NA | 1 | 0 | |
| Kj_022 | Komonfugu | SJ | Shimonoseki, Yamaguchi Pref. | 2006-06-18 | 0 | NA | NA | 1 | 0 | |
| Kj_023 | Komonfugu | SJ | Shimonoseki, Yamaguchi Pref. | 2001-11-20 | 0 | NA | NA | 1 | 0 | |
| Kj_025 | Komonfugu | SJ | Noto, Ishikawa Pref. | 2000-09-19 | 0 | NA | NA | 1 | 0 | |
| Kj_026 | Komonfugu | SJ | Sado Island, Niigata Pref. | 2006-11 | 1 | 0.33507 | 0.46441 | 1 | 1 | SAMD01606017 |
| Kj_027 | Komonfugu | SJ | Sado Island, Niigata Pref. | 2006-11 | 1 | 0.37585 | 0.48486 | 1 | 0 | |
| Kj_066 | Komonfugu | SJ | Kasumi, Hyogo Pref. | 2012-05-26 | 1 | 0.41264 | 0.49133 | 1 | 1 | SAMD01606018 |
| Kj_067 | Komonfugu | SJ | Kasumi, Hyogo Pref. | 2012-06-28 | 1 | 0.30840 | 0.56024 | 1 | 1 | SAMD01606019 |
| Kj_068 | Komonfugu | SJ | Kasumi, Hyogo Pref. | 2012-06-28 | 1 | 0.33938 | 0.31547 | 1 | 0 | |
| Kj_094 | Komonfugu | SJ | Sado Island, Niigata Pref. | 2013-06-05 | 1 | 0.26557 | 0.38545 | 1 | 1 | SAMD01606020 |
| Kj_095 | Komonfugu | SJ | Sado Island, Niigata Pref. | 2013-06-05 | 1 | 0.36737 | 0.53566 | 1 | 0 | |
| Kj_105 | Komonfugu | SJ | Sado Island, Niigata Pref. | 2013-06-05 | 1 | 0.27596 | 0.37426 | 1 | 0 | |
| Kj_137 | Komonfugu | SJ | Kasumi, Hyogo Pref. | 2014-11-27 | 1 | 0.35868 | 0.40148 | 1 | 0 | |
| Kj_138 | Komonfugu | SJ | Kasumi, Hyogo Pref. | 2014-11-27 | 1 | 0.27968 | 0.38142 | 1 | 0 | |
| Kj_139 | Komonfugu | SJ | Kasumi, Hyogo Pref. | 2014-11-27 | 1 | 0.26623 | 0.32911 | 1 | 1 | SAMD01606021 |
| Kj_140 | Komonfugu | SJ | Kasumi, Hyogo Pref. | 2014-11-27 | 1 | 0.29945 | 0.40922 | 1 | 0 | |
| Kj_141 | Komonfugu | SJ | Kasumi, Hyogo Pref. | 2014-11-27 | 1 | 0.31324 | 0.46638 | 1 | 0 | |
| Kj_166 | Komonfugu | SJ | Sado Island, Niigata Pref. | 2015-06-03 | 1 | 0.24164 | 0.33579 | 1 | 1 | SAMD01606022 |
| Kj_167 | Komonfugu | SJ | Sado Island, Niigata Pref. | 2015-06-03 | 1 | 0.30154 | 0.39637 | 1 | 0 | |
| Kj_170 | Komonfugu | SJ | Sado Island, Niigata Pref. | 2016-06-08 | 1 | 0.24975 | 0.45372 | 1 | 0 | |
| Kj_171 | Komonfugu | SJ | Sado Island, Niigata Pref. | 2016-06-08 | 1 | 0.31716 | 0.46263 | 1 | 0 | |
| Kj_172 | Komonfugu | SJ | Sado Island, Niigata Pref. | 2016-06-08 | 1 | 0.25592 | 0.45525 | 1 | 0 | |
| Kj_173 | Komonfugu | SJ | Sado Island, Niigata Pref. | 2016-06-08 | 1 | 0.31797 | 0.43912 | 1 | 0 | |
| Kp_018 | Komonfugu | PO | Minamiise, Mie Pref. | 2006-05 | 1 | 0.33211 | 0.33792 | 1 | 0 | |
| Kp_019 | Komonfugu | PO | Minamiise, Mie Pref. | 2006-05 | 1 | 0.30167 | 0.45874 | 1 | 1 | SAMD01606023 |
| Kp_024 | Komonfugu | PO | Kosai, Shizuoka Pref. | 2000-04-26 | 0 | NA | NA | 1 | 0 | |
| Kp_106 | Komonfugu | PO | Minamiise, Mie Pref. | 2014-01 | 0 | NA | NA | 1 | 0 | |
| Kp_111 | Komonfugu | PO | Minamiise, Mie Pref. | 2014-01 | 1 | 0.41895 | 0.42022 | 1 | 0 | |
| Kp_136 | Komonfugu | PO | Atsumi Peninsula, Aichi Pref. | 2014-03 | 1 | 0.32450 | 0.30112 | 1 | 1 | SAMD01606024 |
| Kp_179 | Komonfugu | PO | Kobe, Hyogo Pref. | 2013-09-18 | 1 | 0.31532 | 0.45372 | 1 | 1 | SAMD01606025 |
| Kp_180 | Komonfugu | PO | Kobe, Hyogo Pref. | 2013-10 | 0 | NA | NA | 1 | 1 | SAMD01606026 |
| Kp_181 | Komonfugu | PO | Minamiise, Mie Pref. | 2014-01 | 0 | NA | NA | 1 | 0 | |
| Kp_182 | Komonfugu | PO | Ashiya, Hyogo Pref. | 2014-11 | 0 | NA | NA | 1 | 1 | SAMD01606027 |
| Kp_183 | Komonfugu | PO | Minamiise, Mie Pref. | 2017-01-04 | 1 | 0.31593 | 0.35152 | 1 | 0 | |
| Kp_184 | Komonfugu | PO | Omaezaki, Shizuoka Pref. | 2017-04-05 | 1 | 0.39978 | 0.54015 | 1 | 0 | |
| Kp_185 | Komonfugu | PO | Omaezaki, Shizuoka Pref. | 2017-04-05 | 1 | 0.38323 | 0.47244 | 1 | 0 | |
| Kp_186 | Komonfugu | PO | Omaezaki, Shizuoka Pref. | 2017-04-05 | 1 | 0.38969 | 0.46152 | 1 | 0 | |
| Kp_187 | Komonfugu | PO | Omaezaki, Shizuoka Pref. | 2017-04-05 | 1 | 0.26907 | 0.41819 | 1 | 0 | |
| Kp_188 | Komonfugu | PO | Omaezaki, Shizuoka Pref. | 2017-04-05 | 1 | 0.33514 | 0.47969 | 1 | 0 | |
| Kp_189 | Komonfugu | PO | Omaezaki, Shizuoka Pref. | 2017-04-05 | 1 | 0.33051 | 0.48599 | 1 | 0 | |
| Kp_190 | Komonfugu | PO | Omaezaki, Shizuoka Pref. | 2017-04-05 | 1 | 0.31140 | 0.29239 | 1 | 1 | SAMD01606028 |
| Kp_191 | Komonfugu | PO | Omaezaki, Shizuoka Pref. | 2017-04-05 | 1 | 0.40901 | 0.45122 | 1 | 0 | |
| Kp_192 | Komonfugu | PO | Omaezaki, Shizuoka Pref. | 2017-04-05 | 1 | 0.35369 | 0.38876 | 1 | 0 | |
| Kp_193 | Komonfugu | PO | Omaezaki, Shizuoka Pref. | 2017-04-05 | 1 | 0.36721 | 0.35492 | 1 | 0 | |
| Kp_194 | Komonfugu | PO | Omaezaki, Shizuoka Pref. | 2017-04-05 | 1 | 0.37241 | 0.42336 | 1 | 1 | SAMD01606029 |
| Kp_195 | Komonfugu | PO | Omaezaki, Shizuoka Pref. | 2017-04-05 | 1 | 0.35863 | 0.38328 | 1 | 0 | |
| Kp_196 | Komonfugu | PO | Omaezaki, Shizuoka Pref. | 2017-04-05 | 1 | 0.35372 | 0.41949 | 1 | 0 | |

| ID | Species | Population | Location | Date | Image | Lightness | Complexity | D-loop | NGS | BioSample |
|---|---|---|---|---|---|---|---|---|---|---|
| Kp_197 | Komonfugu | PO | Omaezaki, Shizuoka Pref. | 2017-04-05 | 1 | 0.35868 | 0.45294 | 1 | 0 | |
| Kp_198 | Komonfugu | PO | Omaezaki, Shizuoka Pref. | 2017-04-05 | 1 | 0.33037 | 0.39701 | 1 | 0 | |
| Kp_199 | Komonfugu | PO | Omaezaki, Shizuoka Pref. | 2017-04-05 | 1 | 0.34869 | 0.48472 | 1 | 0 | |
| Kp_200 | Komonfugu | PO | Omaezaki, Shizuoka Pref. | 2017-04-05 | 1 | 0.33672 | 0.44439 | 1 | 0 | |
| Kp_201 | Komonfugu | PO | Omaezaki, Shizuoka Pref. | 2017-04-05 | 1 | 0.31477 | 0.41203 | 1 | 0 | |
| Kp_202 | Komonfugu | PO | Omaezaki, Shizuoka Pref. | 2017-04-05 | 1 | 0.35426 | 0.46591 | 1 | 0 | |
| Kp_203 | Komonfugu | PO | Omaezaki, Shizuoka Pref. | 2017-04-05 | 1 | 0.30777 | 0.47921 | 1 | 0 | |
| Kp_221 | Komonfugu | PO | Akashi, Hyogo Pref. | 2022-10-20 | 1 | 0.28948 | 0.31914 | 0 | 0 | |
| Kp_222 | Komonfugu | PO | Akashi, Hyogo Pref. | 2022-10-20 | 1 | 0.33017 | 0.40034 | 0 | 0 | |
| Kp_223 | Komonfugu | PO | Akashi, Hyogo Pref. | 2022-10-20 | 1 | 0.32784 | 0.57815 | 0 | 0 | |
| Kp_224 | Komonfugu | PO | Akashi, Hyogo Pref. | 2022-10-20 | 1 | 0.32590 | 0.42067 | 0 | 0 | |
| Kp_225 | Komonfugu | PO | Akashi, Hyogo Pref. | 2022-10-20 | 1 | 0.34308 | 0.42376 | 0 | 0 | |
| Kp_226 | Komonfugu | PO | Akashi, Hyogo Pref. | 2022-10-20 | 1 | 0.32073 | 0.41930 | 0 | 0 | |
| Kp_227 | Komonfugu | PO | Akashi, Hyogo Pref. | 2022-10-20 | 1 | 0.33900 | 0.50682 | 0 | 0 | |
| Kp_228 | Komonfugu | PO | Akashi, Hyogo Pref. | 2022-10-22 | 1 | 0.37175 | 0.44518 | 0 | 0 | |
| Kp_229 | Komonfugu | PO | Akashi, Hyogo Pref. | 2022-10-22 | 1 | 0.35913 | 0.49465 | 0 | 0 | |
| Kp_230 | Komonfugu | PO | Akashi, Hyogo Pref. | 2022-10-22 | 1 | 0.33077 | 0.43145 | 0 | 0 | |
| Kp_231 | Komonfugu | PO | Akashi, Hyogo Pref. | 2022-10-22 | 1 | 0.23390 | 0.39353 | 0 | 0 | |
| Kp_232 | Komonfugu | PO | Akashi, Hyogo Pref. | 2022-10-22 | 1 | 0.35656 | 0.41060 | 0 | 0 | |
| Kp_233 | Komonfugu | PO | Akashi, Hyogo Pref. | 2022-10-22 | 1 | 0.30452 | 0.37941 | 0 | 0 | |
| Kp_234 | Komonfugu | PO | Akashi, Hyogo Pref. | 2022-10-22 | 1 | 0.30319 | 0.37833 | 0 | 0 | |
| Kp_235 | Komonfugu | PO | Akashi, Hyogo Pref. | 2022-10-22 | 1 | 0.38216 | 0.47595 | 0 | 0 | |
| Kp_236 | Komonfugu | PO | Akashi, Hyogo Pref. | 2022-10-22 | 1 | 0.35586 | 0.48576 | 0 | 0 | |

**Table S2: Genome-wide nucleotide diversity (π) of mushifugu and komonfugu.**

Genome-wide nucleotide diversity (π) was calculated from whole-genome resequencing data. Values are shown for each species and for populations from the Sea of Japan (Mj, Kj) and the Pacific Ocean (Mp, Kp).

| Species/Population | π |
|---|---|
| Mushifugu | 0.00212 |
| Mj | 0.00187 |
| Mp | 0.00187 |
| Komonfugu | 0.00324 |
| Kj | 0.00317 |
| Kp | 0.00327 |

**Table S3: Genome-wide absolute divergence ($d_{XY}$) and genetic differentiation ($F_{ST}$) among populations of mushifugu and komonfugu.**

Pairwise genome-wide estimates of absolute nucleotide divergence ($d_{XY}$) and genetic differentiation between populations were calculated using pixy. $F_{ST}$ values are shown as Hudson's estimator (Fst_Hudson_gw) and Weir & Cockerham's weighted estimator (Fst_WC_weighted).

| Pop1 | Pop2 | Dxy_gw | Fst_Hudson_gw | Fst_WC_weighted |
|---|---|---|---|---|
| Mushifugu | Komonfugu | 0.00452 | 0.4073 | 0.3831 |
| Mj | Mp | 0.00235 | 0.2038 | 0.2063 |
| Kj | Kp | 0.00327 | 0.0150 | 0.0147 |
| Mj | Kj | 0.00452 | 0.4428 | 0.4199 |
| Mp | Kp | 0.00453 | 0.4321 | 0.4091 |
| Mj | Kp | 0.00452 | 0.4310 | 0.4082 |
| Mp | Kj | 0.00453 | 0.4443 | 0.4212 |

**Table S4: ABBA–BABA test (D-statistics) for introgression among mushifugu and komonfugu populations.** Results of ABBA–BABA tests conducted using Dsuite. Columns show the test population assignments (P1, P2, P3), the D-statistic, Z-score, and associated p-value, as well as the counts of BBAA, ABBA, and BABA site patterns. Significant excess of allele sharing (|Z| > 3) indicates signals of introgression.

| P1 | P2 | P3 | D-statistic | Z-score | p-value | BBAA | ABBA | BABA |
|---|---|---|---|---|---|---|---|---|
| Mp | Mj | Komonfugu | 0.00575 | 2.666 | 0.0077 | 388821 | 70531.2 | 69724.4 |
| Mp | Mj | Kj | 0.00753 | 3.166 | 0.0015 | 388294 | 70548 | 69493.3 |
| Mp | Mj | Kp | 0.00416 | 2.013 | 0.0441 | 388250 | 70243.2 | 69661.2 |
| Kp | Kj | Mushifugu | 0.00044 | 0.210 | 0.8339 | 309790 | 102352 | 102261 |
| Kp | Kj | Mj | 0.00164 | 0.731 | 0.4650 | 308583 | 102170 | 101836 |
| Kj | Kp | Mp | 0.00073 | 0.359 | 0.7193 | 309389 | 102016 | 101868 |

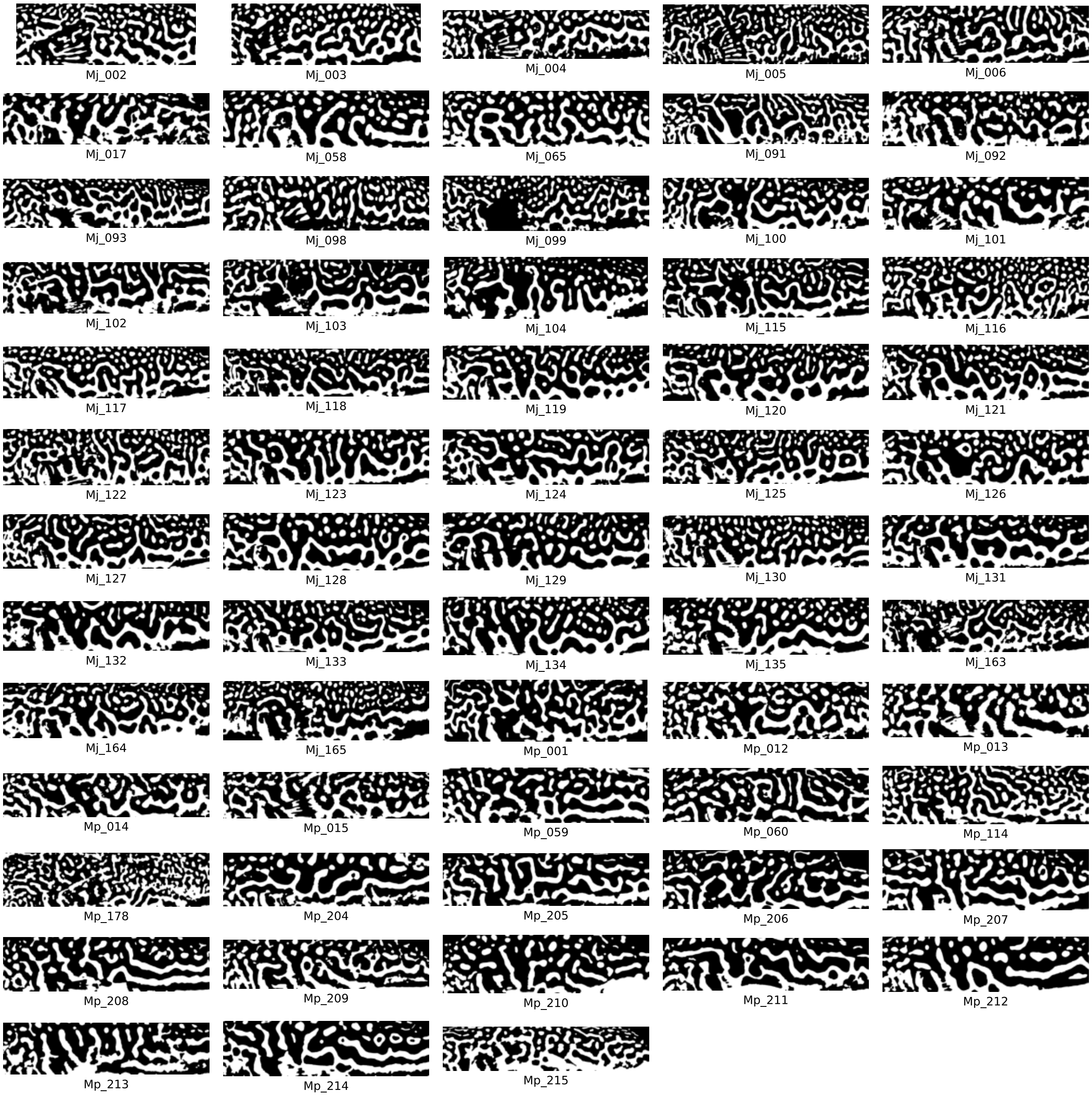


**Supplementary Figure S1: Body patterns of mushifugu (*T. exascurus*).**

Binarized images used for quantitateve analysis of body pattern complexity.

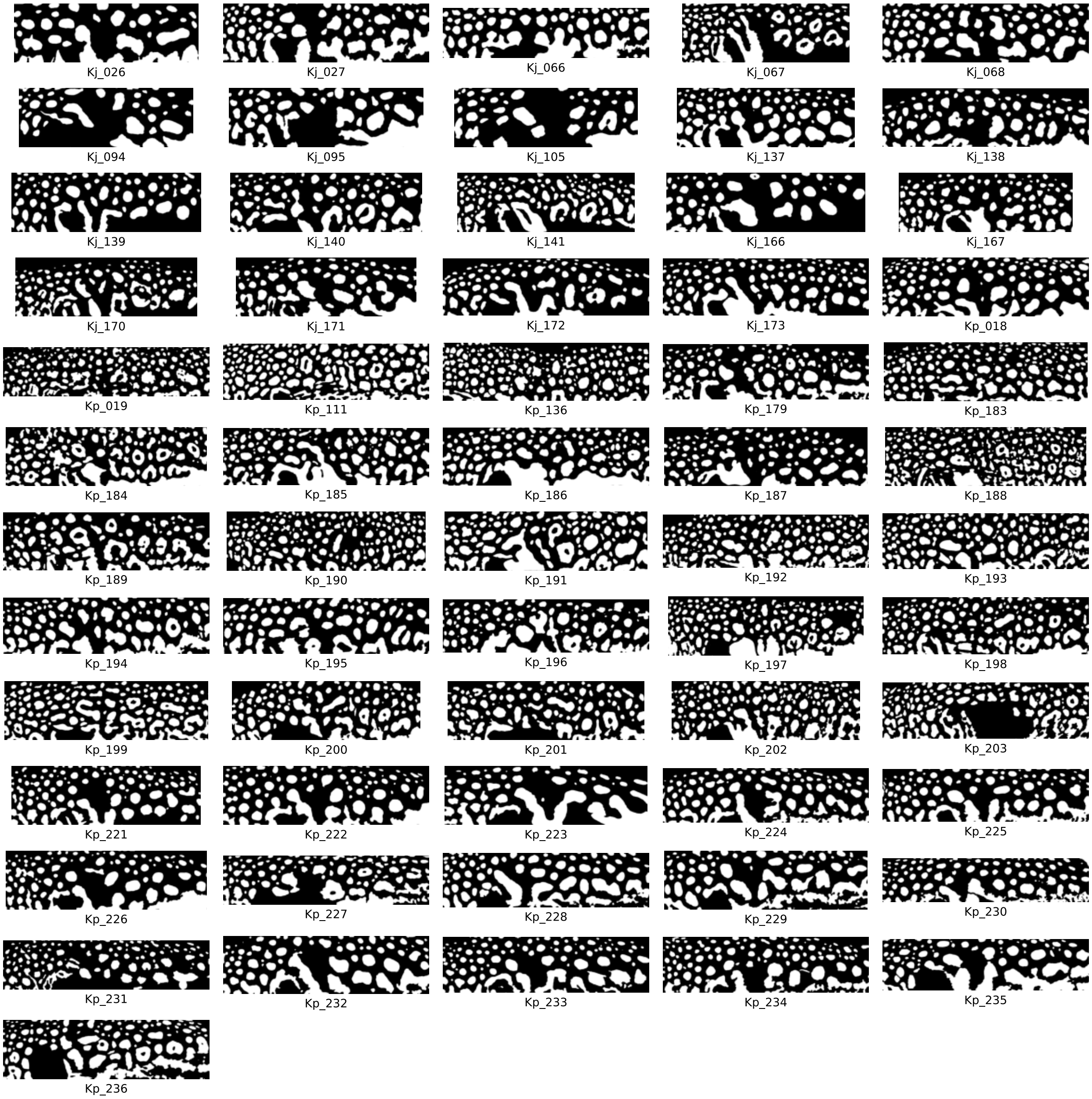


**Supplementary Figure S2: Body patterns of komonfugu (*T. flavipterus*).**

Binarized images used for quantitateve analysis of pattern complexity.

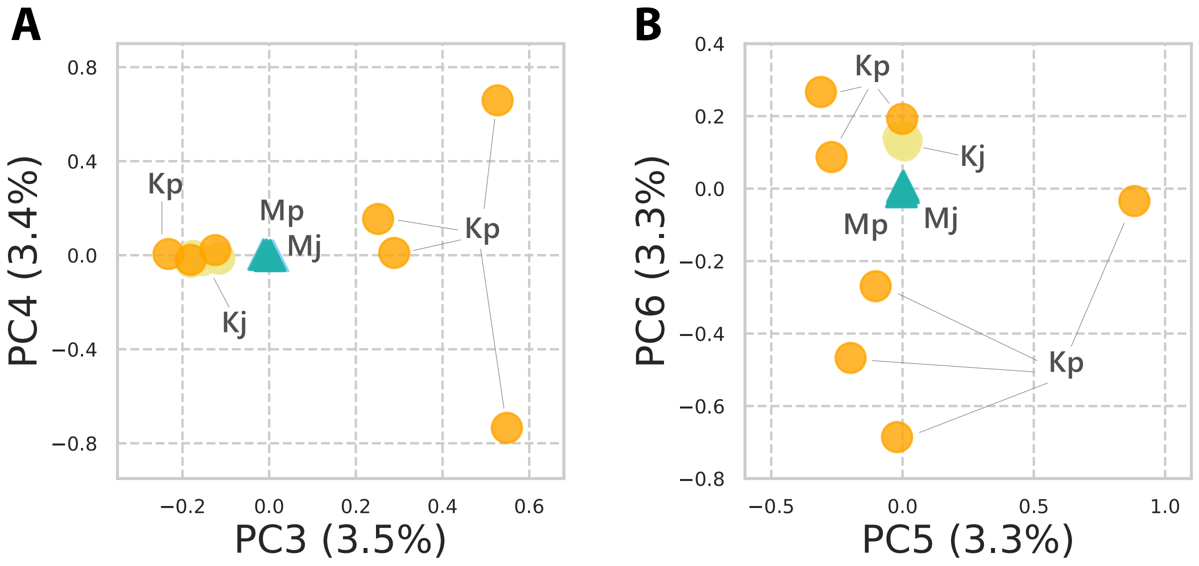


**Supplementary Figure S3: Principal component analysis (PCA) of genome-wide SNP variation.**

(A) Distribution of samples along the third and fourth principal components (PC3, 3.5%; PC4, 3.4%). (B) Distribution along the fifth and sixth principal components (PC5, 3.3%; PC6, 3.3%). Abbreviations: Mj, mushifugu (Sea of Japan); Mp, mushifugu (Pacific coast); Kj, komonfugu (Sea of Japan); Kp, komonfugu (Pacific coast).
